# Linking Muscle Mechanics to the Metabolic Cost of Human Hopping

**DOI:** 10.1101/2023.01.31.526545

**Authors:** Luke N. Jessup, Luke A. Kelly, Andrew G. Cresswell, Glen A. Lichtwark

**Affiliations:** School of Human Movement and Nutrition Sciences, Centre for Sensorimotor Performance, The University of Queensland, Brisbane, QLD, Australia

**Keywords:** Biomechanics, Muscle, Energetics, Locomotion, Hopping

## Abstract

Many models have been developed to predict metabolic energy expenditure based on biomechanical proxies of muscle function. However, current models may only perform well for select forms of locomotion, not only because the models are rarely rigorously tested across subtle and broad changes in locomotor task, but also because previous research has not adequately characterised different forms of locomotion to account for the potential variability in muscle function and thus metabolic energy expenditure. To help to address the latter point, the present study imposed frequency and height constraints to hopping and quantified gross metabolic power as well as the activation requirements of medial gastrocnemius, lateral gastrocnemius (GL), soleus (SOL), tibialis anterior, vastus lateralis (VL), rectus femoris (RF) and biceps femoris (BF), and the work requirements GL, SOL and VL. Gross metabolic power increased with a decrease in hop frequency and increase in hop height. There was no hop frequency or hop height effect on the mean electromyography (EMG) of ankle musculature, however, the mean EMG of VL and RF increased with a decrease in hop frequency and that of BF increased with an increase in hop height. With a reduction in hop frequency, GL, SOL and VL fascicle shortening, fascicle shortening velocity and fascicle to MTU shortening ratio increased, whereas with an increase in hop height, only SOL fascicle shortening velocity increased. Therefore, within the constraints that we imposed, decreases in hop frequency and increases in hop height resulted in increases in metabolic power that could be explained by increases in the activation requirements of knee musculature and/or increases in the work requirements of both knee and ankle musculature.

**Summary Statement:** This study directly measures activation and work requirements of lower-limb musculature and whole-body metabolic energy requirements across a wide variety of human hopping conditions, helping to guide biomechanical models of energy expenditure.

## Introduction

Mapping the mechanistic link between locomotor mechanics and energetics has proven to be difficult. Many models have been developed to predict metabolic energy expenditure based on biomechanical proxies of muscle function. Some suggest derivatives of mechanical work as predictors of energy expenditure (e.g., Cavagna and Kanecko, 1977; Donelan et al., 2002; Riddick and Kuo, 2022; Taylor and Heglund, 1982; Willems et al., 1995), some suggest derivatives of force (e.g., Chasiotis et al., 1987; Kipp et al., 2018; Kram and Taylor, 1990; McMahon et al., 1987), while others propose a combination of the two (e.g., Alexander, 2002; Pontzer, 2016). However, current models are rarely rigorously tested across subtle and broad changes in locomotor task and may be biased to select forms of movement.

The variable energetic cost of human hopping across different hop frequency-height combinations provides a good example of the difficulty in predicting energy expenditure based on mechanics. Gutmann and Bertram (2013) measured metabolic energy expenditure during vertical hopping in humans while broadly constraining hop frequency (from ~ 1.4 – 3.2 hops s-1) and hop height (from ~ 0.06 to 0.22 m). They developed extensive cost landscapes across conditions and observed an increase in metabolic power at both lower frequencies and higher heights. Gutmann and Bertram (2017a, b) also tested a range of cost models on this hopping data and found that while muscle impulse- and work-based models outperformed muscle mean-force- and force-rate-based models for predicting energy expenditure, a considerable amount of variance was unexplained in all models. Unfortunately, these analyses were limited to using ground reaction force data, and as such, were unable to directly quantify how muscle function changes with hop frequency, hop height, and metabolic power.

Whilst we know a considerable amount about the mechanisms that drive energy expenditure in muscles during contraction (Barclay, 2023; Brooks, 2012; Woledge et al., 1985), direct measurements of muscle mechanics are essential for understanding muscle energetics during gross movements because of variability that can be attributed to the elastic function of in-series tendinous tissue. Movements that more effectively utilise tendon to recycle the work that is done upon muscle by body weight can minimise muscle activation and work requirements, which is energetically favourable (Fukunaga et al., 2001; Hof et al., 2002; Lichtwark and Wilson 2005a,b; Roberts et al., 1997). Specifically, the function of tendon can reduce muscle activation by reducing the length change and velocity requirements of muscle contractions. Furthermore, reducing muscle shortening velocity can also further reduce the required metabolic rate associated with work production (known as the Fenn effect (Barclay, 2023)).

Studies have shown that the interaction between muscle and tendon is highly sensitive to constraints on movement tasks. For instance, Dean and Kuo (2011) measured muscle activation and metabolic energy expenditure, and estimated length changes of calf muscle fascicles, while constraining the frequency at which participants performed a cyclical bouncing task. Consistent with other frequency-constrained studies on gait (Brennan et al., 2017; Högberg, 1952; Ishikawa et al., 2005; Monte et al., 2021; Ralston, 1958; Robertson and Sawicki, 2015; Swinnen et al., 2022; Takeshita et al., 2006; Waugh et al., 2017), the authors observed a U-shaped relationship between movement frequency and metabolic power that could be explained by muscle mechanics. Participants became increasingly ‘tuned’ to exploit tendon compliance and minimise active muscle work the closer they moved to an ‘energetically ideal’ frequency. Although, as participants moved further below or above this frequency, temporal shifts occurred between muscle and tendon dynamics that caused muscle to do more active work. These findings can also be contextualised in terms of a spring-mass system operating at a resonant frequency to which the system is optimally tuned, and that moving above or below that frequency requires additional energy (e.g., from muscle) to drive the system (Robertson and Sawicki, 2014).

Few studies have made direct, concurrent measures of muscle activation, fascicle dynamics, and metabolic energy expenditure across movement tasks. Beck et al. (2020) found that decreasing the duty factor during cyclical ankle tasks (i.e., producing the same peak plantar flexor moment over a shorter duration) increased metabolic power by amplifying peak muscle-tendon forces, thereby increasing fascicle shortening and shortening velocity through increased muscle activation. Van der Zee and Kuo (2021) found that increasing movement frequency during cyclical knee tasks increased metabolic power in a way that they argued was more consistent with amplified muscle activation (i.e., ion transport) than muscle work (i.e., cross-bridge cycling). However, they did not report fascicle shortening velocities, which may have contributed to changes in both activation and metabolic rate (Fletcher et al., 2013; Lichtwark and Wilson, 2008).

Although the aforementioned studies were performed on single joints with a single movement constraint, their findings provide compelling mechanistic rationale for why energy expenditure may vary between movements. However, as we scale up to locomotion, movements are less likely to be characterised by a single constraint (e.g., frequency), meaning that these studies cannot account for all potential variability in muscle function and thus metabolic energy expenditure.

The present study was designed to impose similar frequency and height constraints to hopping as Gutmann and Bertram (2013) in order to illustrate how changes in hopping parameters relate to changes in activation and work requirements of major muscle groups in the lower limb. This was done in order to better characterise how different conditions affect energy expenditure with respect to the requirements of the movement, and to use these findings to help to direct future biomechanical models of energy expenditure. We hypothesised that metabolic power would have a moderate positive relationship with hop height and a moderate negative relationship with hop frequency, within the constraints imposed. We also hypothesised that increases in metabolic power would be reflected by muscle mechanics that are less energetically favourable, as seen by increases in muscle activation, fascicle shortening, fascicle shortening velocity, and fascicle to muscle-tendon unit shortening ratio.

## Materials and Methods

### Ethics

This study was approved by, and conducted in accordance with, the University of Queensland Health and Behavioural Sciences, Low and Negligible Risk Ethics Sub-Committee (2020002026).

### Participants

Twelve healthy participants (8 male, 4 female), age 26 ± 4 (mean ± SD) years, height 173 ± 10 cm, and mass 69 ± 15 kg provided written consent to participate in this study. All participants regularly engaged in strenuous aerobic exercise (i.e., ≥ 30 min, three times per week of running, cycling, football etc.).

### Protocol

Participants were directed to hop in place on both feet with arms held rigidly at their side – any other observed movement which would otherwise complicate the interpretation of the results was limited.

Participants hopped for 4 min per trial. Metabolic energy consumption was recorded throughout each trial. Kinetics, kinematics, and muscle mechanics were recorded simultaneously for 5 sec at the 3 min mark of each trial. Nineteen trials split across four testing sessions, were performed by each participant. Each trial was separated by a minimum of 5 min rest and until subjects felt fully recovered. The four sessions were spaced by a minimum of seven days. Trials were either unconstrained, height-constrained, frequency-constrained, or fully constrained (Fig. 1). All height and frequency constraints were chosen to be in a similar range to those reported by Gutmann and Bertram (2013), and were pilot tested to ensure that participants could undertake them for 4 min.

**Figure 1.**
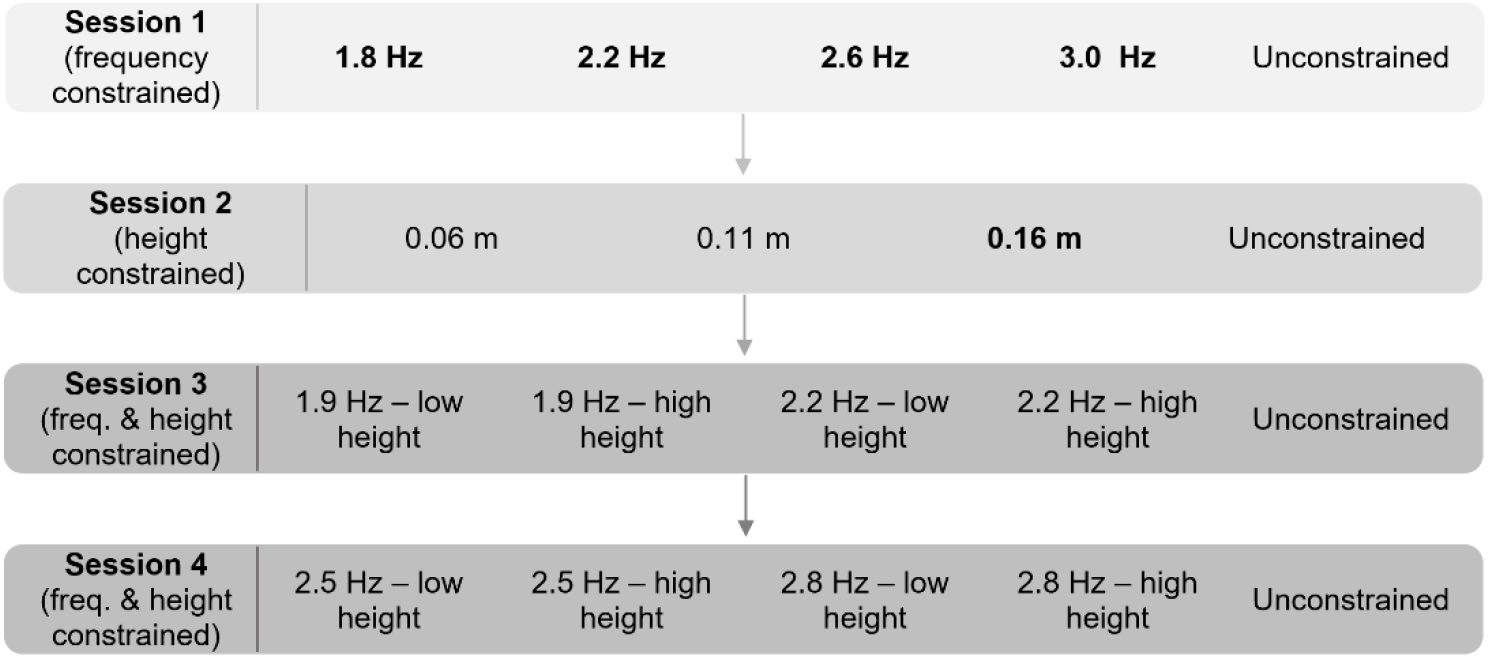
Example layout of the four testing sessions. All of the conditions that were prescribed across the four testing sessions. The conditions that are bolded were those used in the group analysis.

For unconstrained trials, participants were free to select their preferred hop frequency and height. One unconstrained trial was performed at random per testing session.

For height-constrained trials, hop height was specified by a bar resting horizontally above the participants’ head, though they were free to select their preferred hop frequency. Three different height-constrained trials – 0.06 m, 0.11 m, and 0.16 m – were performed at random during one of the testing sessions.

For frequency-constrained trials, hop frequency was specified by a metronome, though participants were free to select their preferred hop height. Four different frequency-constrained trials – 1.8 Hz, 2.2 Hz, 2.6 Hz, and 3.0 Hz – were performed at random during one of the testing sessions.

For fully constrained trials, both hop height and frequency were set as above. Four different hop frequencies – 1.9 Hz, 2.2 Hz, 2.5 Hz, and 2.8 Hz – were used in tandem with both a low and high hop height for a total of eight different height-frequency combinations. To determine the heights for each frequency, participants were instructed to hop as low and as high as possible at each frequency for ~ 20 sec, twice. These heights were measured by manually adjusting the bar above the participants’ head to the height that they were hopping. The heights were then revised and set based on what we deemed would ensure steady-state, aerobic respiration for the ensuing 4 min trials. There was a mean difference of 0.05 ± 0.01 m between the fully constrained low height and fully constrained high height conditions. The eight fully constrained trials were randomly split between the last two testing sessions.

Participants were provided with verbal feedback throughout each trial to maximise adherence to the constraints.

### Kinetics and Kinematics

Ground reaction force (GRF) data was collected at 1000 Hz from two floor embedded force plates (OR6-7, AMTI, MA, USA). An eleven-camera optoelectronic motion capture system (Oqus, Qualisys, AB, Sweden) was used to capture the position of 43 reflective markers at 200 Hz and was time-synchronised with the GRF data. Markers were placed on anatomical landmarks: both legs at the distal phalanx of the first toe, metatarsophalangeal joints 1 and 5, posterior process of the calcaneus, medial and lateral malleolus of the ankle, lateral mid-shank, medial and lateral joint rotation centre of the knee, and lateral mid-thigh. Markers were also placed on the left and right anterior superior iliac spine, posterior superior iliac spine, and iliac crest directly superior to the greater trochanter, as well as on the coccyx, suprasternal notch of the manubrium, C7 spinous process, and acromion process of the left and right shoulder. Raw marker trajectories were filtered using a 5-point moving average using Qualisys Track Manager software. All data were digitally exported for analysis in OpenSim (Delp et al., 2007). An OpenSim model previously described by Lai et al. (2017) was applied to the data. Scaling factors were calculated by dividing participant-specific marker distances by the corresponding distances on the generic model. Body kinematics during hopping trials were determined using OpenSim’s inverse kinematics tool, which uses a weighted least-squares fit of the generic markers rigidly attached to the scaled model to the experimental markers. Body kinetics were determined using OpenSim’s inverse dynamics tool – inverse kinematics and GRF data were filtered at 15 Hz prior to inverse dynamic calculation (Kristianslund et al., 2012). Inverse kinematic and dynamic results for each trial were imported into MATLAB (MathWorks, MA, USA) where a custom script differentiated joint angles with respect to time to determine joint velocities. These were subsequently multiplied by joint moments to calculate joint powers, and then integrated over time to calculate joint work. Vertical centre of mass (CoM) power was determined by first integrating CoM accelerations calculated from GRF data to obtain CoM velocities and then multiplying CoM velocities by the GRF data, as per the method outlined by Smith et al. (2021). Time-series data were cropped from ground contact through to the end of the subsequent flight phase for each hop (events indexed based on GRF data). Hop frequency was calculated as 1 / hop duration (s). Hop height was calculated as the difference between the vertical coccyx marker position at the top of flight and that during the static, standing measurement. All kinematic and kinetic data, as well as muscle data, were averaged across five hop cycles per trial.

### Muscle-Tendon Mechanics

B-mode ultrasound (ArtUs EXT-1H, Telemed, Vilnius, Lithuania) was used to visualise muscle fascicle lengths in-vivo (Lichtwark and Wilson, 2006). Specifically, a linear array transducer (LV8-5N60-A2, Telemed, Vilnius, Lithuania) was used to image lateral gastrocnemius (GL) and soleus (SOL) muscle fascicles, and a second transducer of the same type was used to image vastus lateralis (VL) muscle fascicles. Both transducers were fixed to the participant using a self-adhesive bandage (Medichill, WA, Australia) such that they maintained longitudinal alignment to the plane of the fascicles throughout each hop (Farris and Lichtwark, 2016). Probe depth was set to 50 mm and recorded using EchoWave II software (Telemed, Vilnius, Lithuania) operated on separate laptop computers. Data from the transducers was synchronised to the motion capture and GRF data, using an external trigger (frame by frame) that captured frames at 160 Hz on both machines. Data was exported to MATLAB, where a tracking software was used to make fascicle length measurements (Cronin et al., 2011; Farris and Lichtwark, 2016). This involved automated tracking using an affine flow algorithm, followed by manual corrections where realignment of a measured fascicle was necessary. To minimise bias, the same fascicle region, detected by an automated software, was tracked for data across all trials per session for each participant (Seynnes and Cronin, 2020). The instantaneous MTU length of GL, SOL and VL was determined by applying OpenSim’s muscle analysis tool to the inverse kinematics results. Fascicle and MTU lengths were then cropped from ground contact through to the end of the subsequent flight phase for each hop. For each participant, fascicle and MTU lengths were normalised to the respective mean fascicle length across all trials. This ensured correct scaling of the MTU lengths for fascicle shortening and the calculation of MTU shortening ratio. Fascicle and MTU length changes were calculated over the period of positive CoM work (Fig. S1) – the most energetically demanding part of the hop cycle since extensor musculature will be actively producing force whilst shortening to some degree. Fascicle velocity was calculated over the same period as the derivative of the normalised length signal.

### Muscle Activations

Surface electromyography (EMG) data was collected to measure the activation of GL, medial gastrocnemius (GM), SOL, tibialis anterior (TA), VL, rectus femoris (RF), and biceps femoris (BF). A ground electrode was positioned on the skin overlying the fibular head. The recording sites were prepared by shaving and then cleaning the skin using an abrasive gel (NuPrep, Weaver and Company, CO, USA) and alcohol. 24 mm diameter EMG electrodes (Covidien, MA, USA) were placed in a bipolar configuration over each muscle, according to SENIAM guidelines (Hermens et al., 1999). Each recording site was marked with an indelible marker. Following each session, participants were encouraged to maintain the marks between sessions, thereby making it easier to place electrodes in the same position between days. EMG signals were amplified by 1000-times, hardware filtered with a bandwidth of 10–2,000 Hz (MA422, Motion Lab Systems, CA, USA) and recorded in Qualisys Track Manager at 2000 Hz. EMG signals were processed in MATLAB where the DC offset was removed using a high-pass filter at 25 Hz, followed by signal rectification, and finally low-pass filtering at 10 Hz to create an EMG envelope (Kirtley, 2006). The EMG signals were then normalised to the maximal activity measured from three maximal squat jumps performed at the beginning of each testing session (Besomi et al., 2020). EMG was then cropped from ground contact through to the end of the subsequent flight phase for each hop, with the mean EMG for each muscle calculated over this period to quantify how active muscles were over time, which is paramount from an energetic standpoint.

### Metabolic Energy Consumption

Oxygen consumption and carbon dioxide elimination was measured with a portable spirometry system (Metamax 3B; Cortex, Leipzig, Germany). The rate of oxygen consumption was required to reach a plateau (i.e., steady-state respiration) within the first 2 min of each trial. To ensure primarily oxidative metabolism, trials were terminated where the respiratory exchange ratio (RER) rose above 1.0 or heart rate above 90% of the age-predicted maximum (0.9 x (220 – age in years)). The effects of diet on metabolic rate were also mitigated by testing 2 h post-prandial and by testing prior to any daily caffeine consumption. Regression analyses were performed in MATLAB on the cumulative integral of the final 2 min of oxygen consumption data of each trial to ensure high linearity (r^2^ > 0.99). Subsequently, the data was averaged and converted to gross metabolic power using standard equations to obtain a representative steady-state value for each trial (Brockway, 1987).

With the amount of rest provided within and between sessions, the effects of fatigue and possible muscle soreness were mitigated. 18 out of 228 total trials were discarded either because they were too aerobically challenging for the participant, and/or because the participant could not adhere to the designated frequency and/or height.

### Statistics

Our preliminary statistical analysis involved determining whether metabolic power (response variable) was positively or negatively related to hop height (fixed effect) and hop frequency (fixed effect) across our entire dataset of 210 trials. To do this, we used linear models (LMs) and the resulting adjusted correlation coefficients (adjusted *R^2^*) to determine the direction and strength of association between each of our fixed effects and the response variable. We also ran a regression model that included both of our fixed effects to test by how much this would strengthen the resulting adjusted *R^2^*. In these LMs, each participant was used as a categorical predictor to account for variation due to participant in the data used in the regression (Bland and Altman, 1995). The formulas used can be written in Wilkinson notation as:

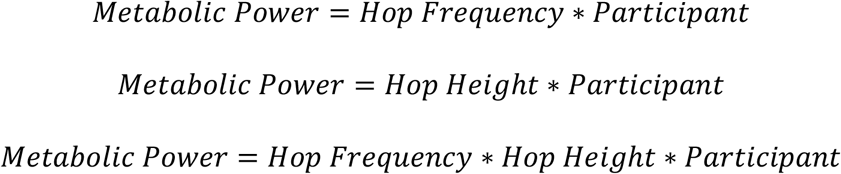

Our next statistical analysis involved comparing metabolic power, muscle fascicle and tendon dynamics, muscle activation, and joint work (response variables) between different groupings of hop height-frequency combinations. The groups selected allowed us to determine whether changes in hop height and hop frequency had an effect on metabolic power and different muscle mechanics, and to visualise whether trends in metabolic power were reflected by trends in particular muscle mechanics. Each group was derived from specific conditions of the data collection (Fig. 1) that each participant performed which forced them to hop in the regions of the frequency-height surface (Fig. 3a) that we wished to compare. Five groups were used as fixed effects in a linear mixed-effects model (LMEM) analysis – low hop frequency and high hop height (LFHH; 1.82 ± 0.09 Hz, 0.19 ± 0.01 m); low hop frequency and low hop height (LFLH; 1.88 ± 0.11 Hz, 0.09 ± 0.02 m); low-medium hop frequency and low hop height (LMFLH; 2.25 ± 0.05 Hz, 0.09 ± 0.02 m); medium-high hop frequency and low hop height (MHFLH; 2.59 ± 0.11 Hz, 0.08 ± 0.02 m); and high hop frequency and low hop height (HFLH; 2.98 ± 0.05 Hz, 0.07 ± 0.01 m). The groups were considered to have a significant effect on the response variable when the *p*-value for the *F*-test on the group fixed-effect coefficient was less than 0.05. LMEMs were used in this case to cope with rare instances of missing data for particular groups for particular response variables, and so that we could treat each participant as a random effect. The formula used can be written in Wilkinson notation as:

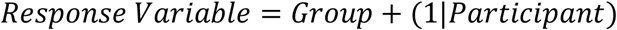

Finally, we wished to further elucidate whether there may have been an effect of hop frequency or hop height on each of our muscle- and joint-based response variables. For each response variable, where there was a significant effect of grouping we ran Sidak multiple comparisons tests to examine whether there was a significant mean difference (Adjusted *p*-value < 0.05) between data from the LFLH versus HFLH group (termed the *frequency effect*) and the LFLH versus LFHH group (termed the *height effect*) – indicating effects of hop frequency and hop height, respectively.

## Results

### Metabolic Power

A moderate positive correlation (*R^2^* = 0.46) was found for hop height and a moderate negative correlation (*R^2^* = 0.41) was found for hop frequency (Fig. 2). When including both hop height and hop frequency as predictors in the regression model, a greater amount of the variance was explained than when considered singularly (*R^2^* = 0.53), however this was still only a moderate to strong correlation. There was a significant effect of grouping on gross metabolic power (p < 0.001). A significant hop frequency (p = 0.003) and hop height (p = 0.013) effect was also found.

**Figure 2.**
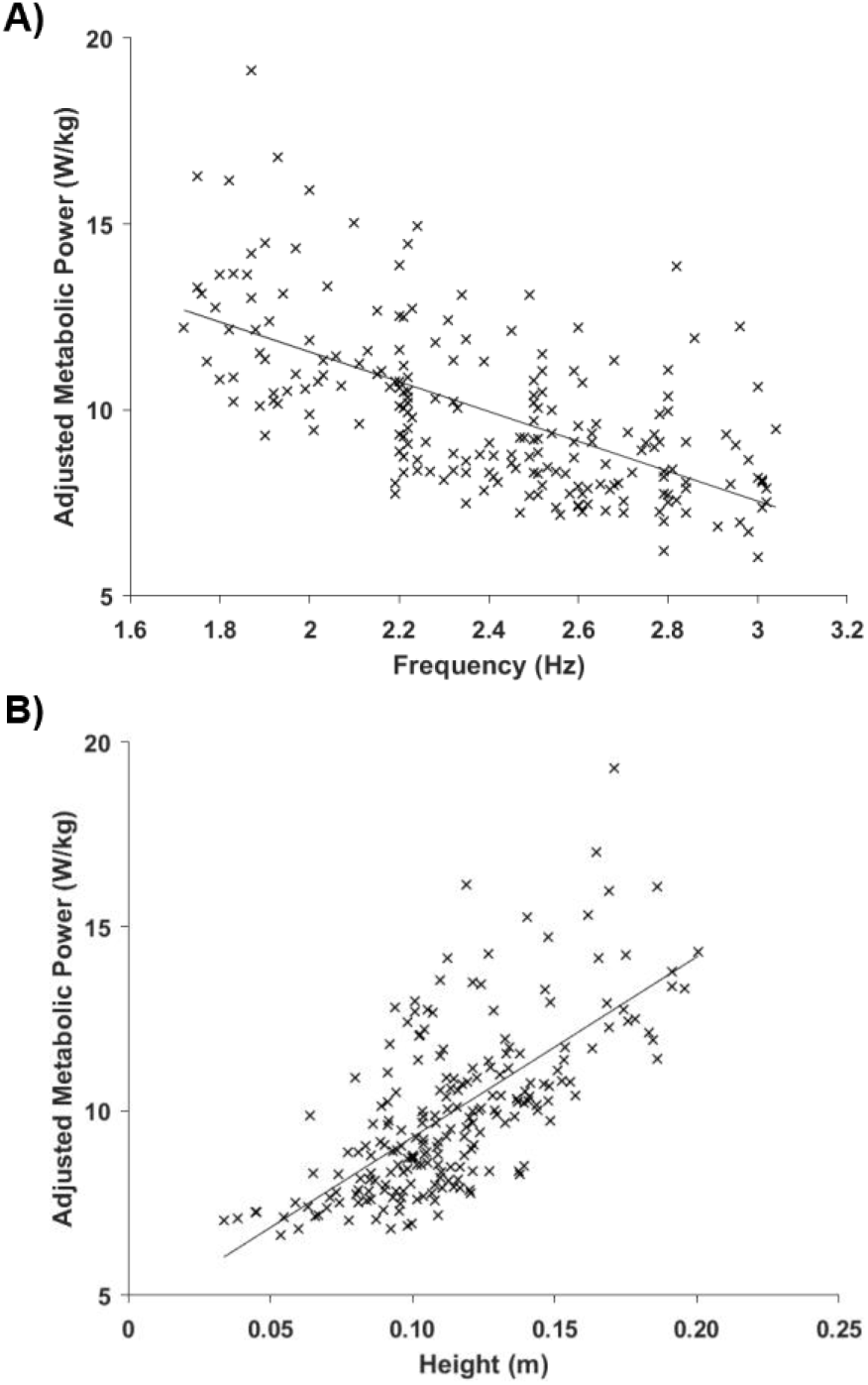
Linear model of gross metabolic power against hop frequency and against hop height. A) Linear model, including regression line, of gross metabolic power (W/kg) against hop frequency (Hz), and B) against hop height (m), across our entire dataset.

**Figure 3.**
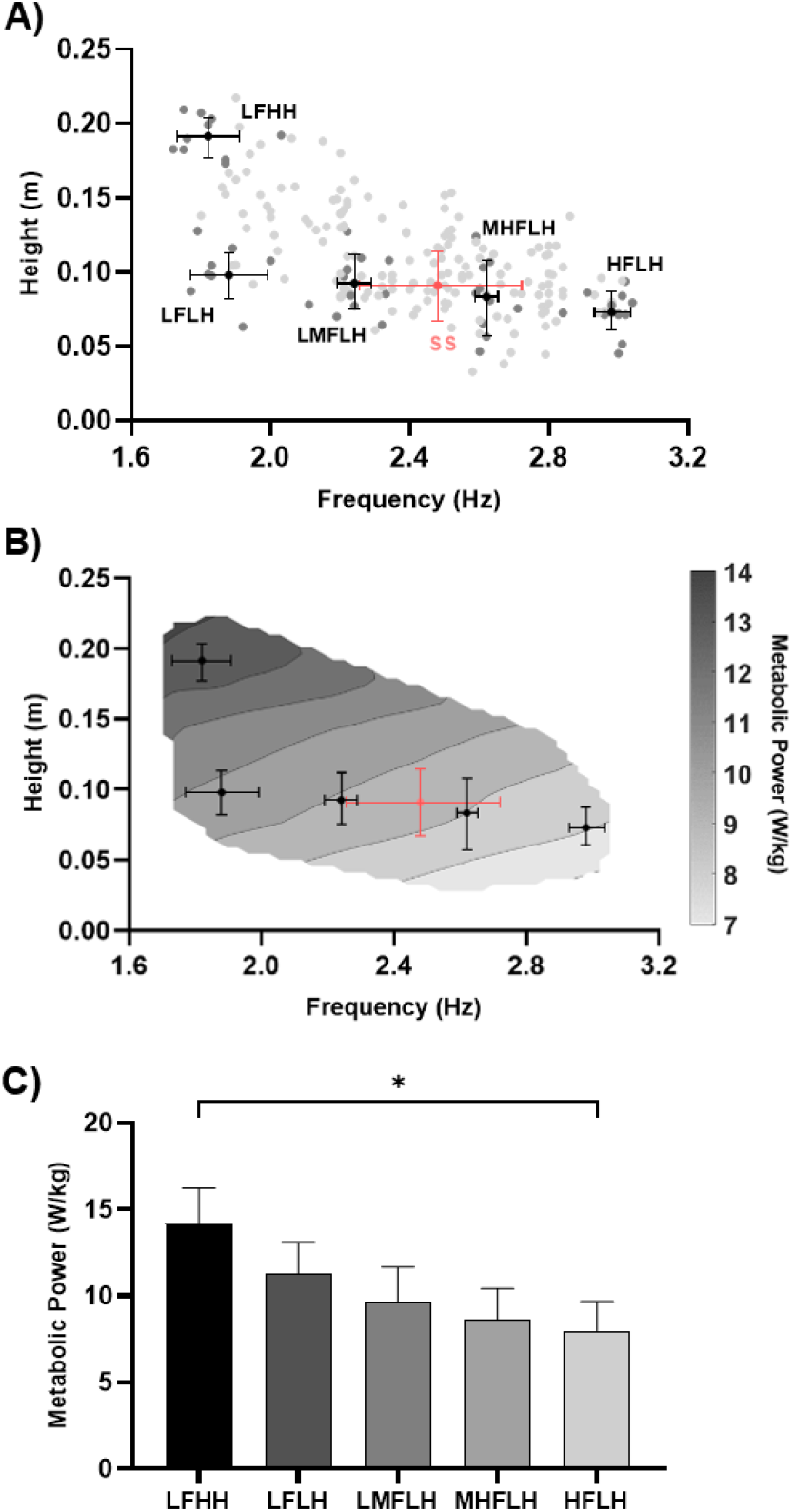
Hop frequency, hop height, and gross metabolic power across the entire dataset and groups of trials. A) The mean ± SD height-frequency of each group analysed (black) and of every self-selected condition performed (salmon), plotted atop the height-frequency of every trial in our dataset (grey), and B) plotted atop the gross metabolic power (W/kg) surface from every trial in our dataset. C) The mean ± SD gross metabolic power (W/kg) for each of the groups analysed. * indicates a significant effect of grouping (p < 0.05).

### Muscle Activation

For muscles crossing the ankle, aside from GM (p = 0.013), there was no significant effect of grouping on mean EMG (SOL (p = 0.845), GL (p = 0.210), TA (p = 0.055)) (Fig. 4). No significant hop frequency (p = 0.548) or hop height (p = 0.227) effect was found for the mean EMG of GM. For muscles crossing the knee, there was a significant effect of grouping on mean EMG for VL, RF, and BF (all p < 0.001). A significant hop frequency effect was found for the mean EMG of VL (p = 0.043) and RF (p = 0.011) but not BF (p = 0.520). A significant hop height effect was found for the mean EMG of BF (p = 0.024) but not VL (p = 0.980) or RF (p = 0.652).

**Figure 4.**
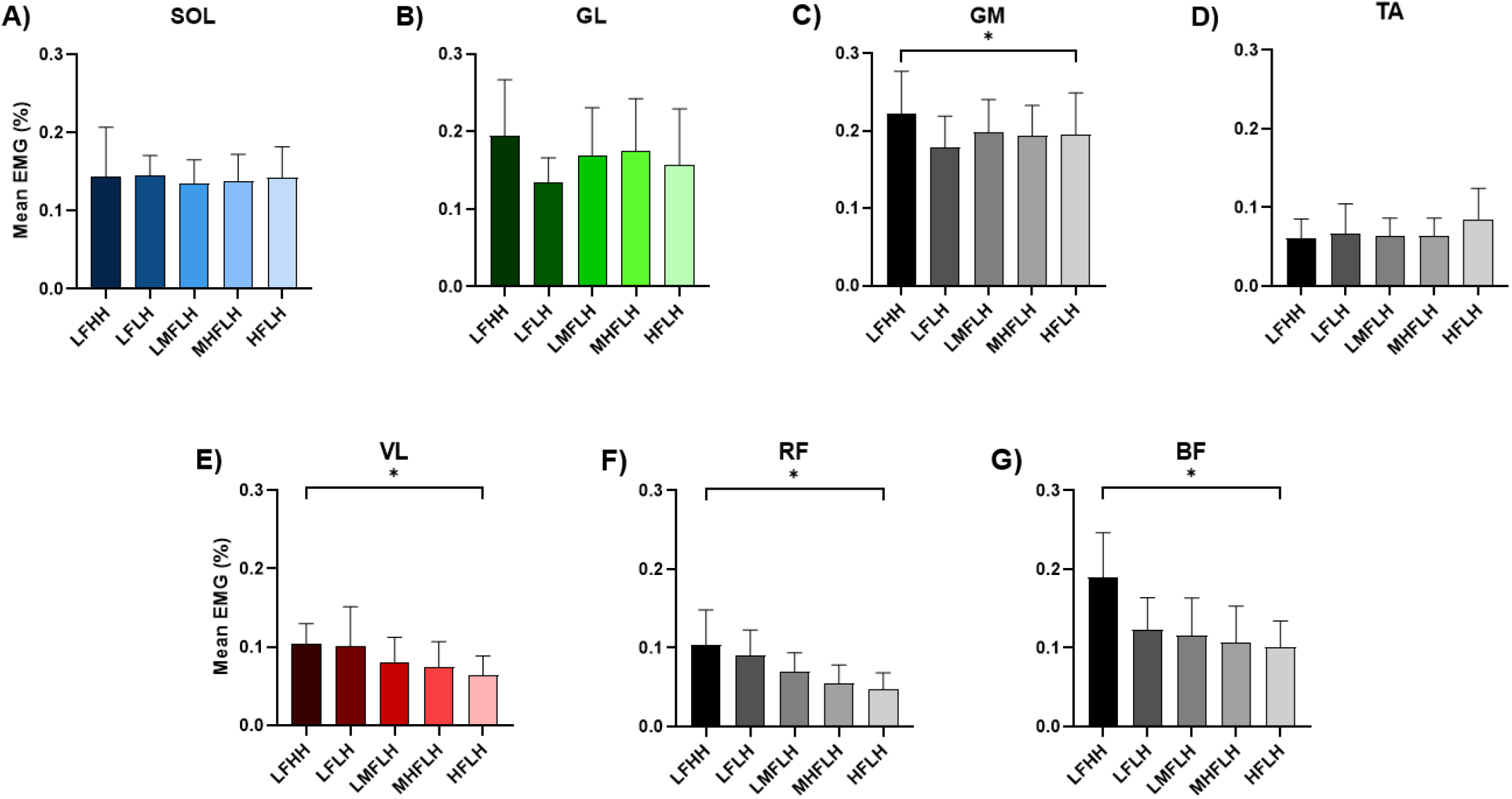
Group analysis of mean EMG data. The mean ± SD mean normalised EMG (%) for each of the groups analysed for SOL (A), GL (B), GM (C), TA (D), VL (E), RF (F), and BF (G). * indicates a significant effect of grouping (p < 0.05).

### Muscle Fascicle and Tendon Dynamics

There was a significant effect of grouping on SOL, GL and VL fascicle shortening, fascicle shortening velocity and fascicle to MTU shortening ratio (all p < 0.001) (Fig. 5). A significant hop frequency effect was found for the fascicle shortening (SOL (p < 0.001), GL (p < 0.001), VL (p = 0.002)), fascicle shortening velocity (SOL (p = 0.027), GL (p < 0.001), VL (p = 0.004)) and fascicle to MTU shortening ratio (SOL (p = 0.005), GL (p = 0.002), VL (p = 0.041)) of all muscles. A significant hop height effect was found for the fascicle shortening velocity of SOL (p = 0.039) but not GL (p = 0.104) or VL (p = 0.919), and no significant hop height effect was found for the fascicle shortening (SOL (p = 0.410), GL (p = 0.740), VL (p = 0.865)) or fascicle to MTU shortening ratio (SOL (p = 0.810), GL (p = 0.367), VL (p = 0.814)) of all muscles.

**Figure 5.**
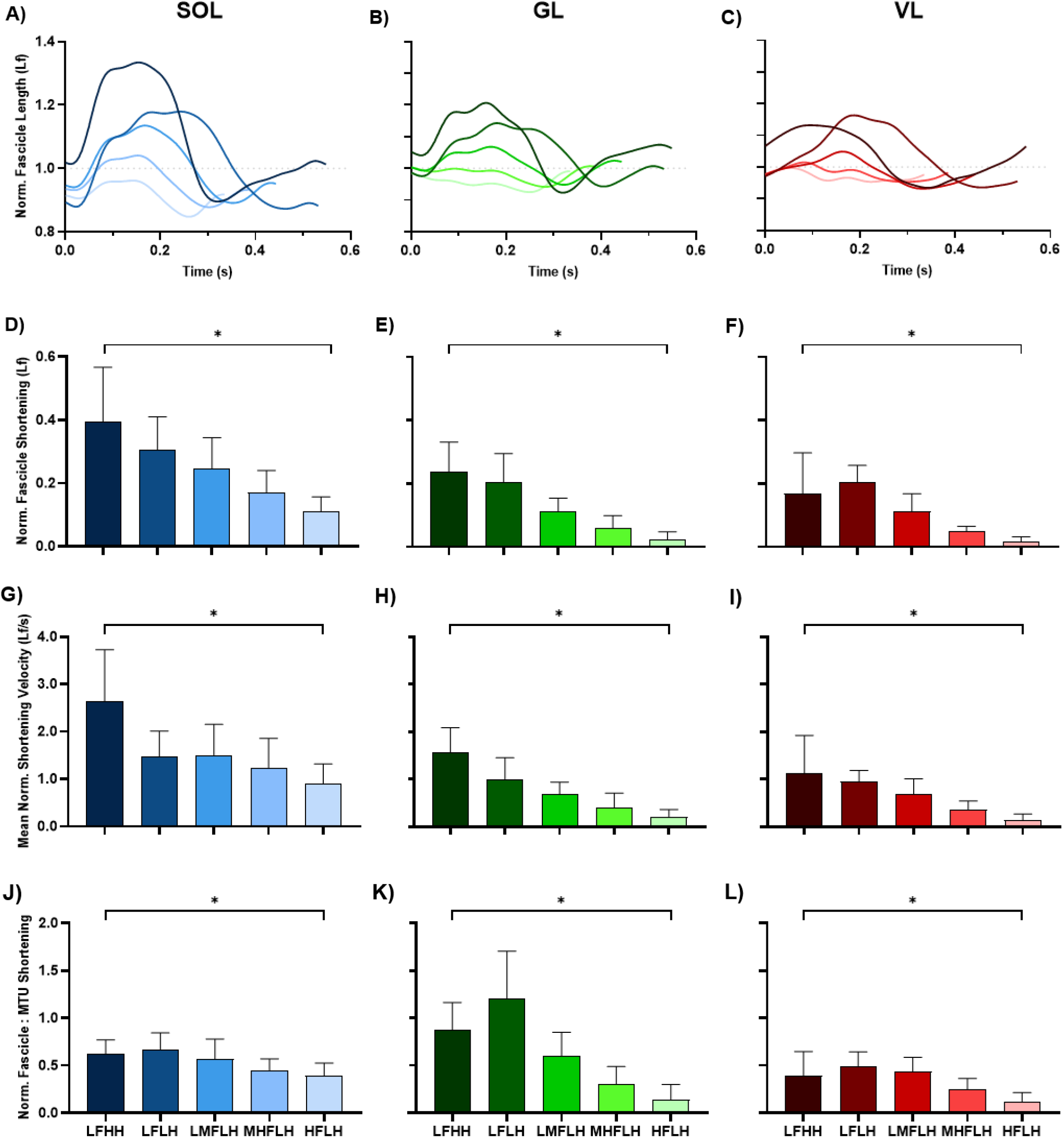
Group analysis of fascicle shortening, fascicle shortening velocity, and fascicle to MTU shortening ratio data. Time-series plots of the mean normalised fascicle length (Lf) for each of the groups analysed for SOL (A), GL (B), and VL (C); and bar plots of the mean ± SD normalised fascicle shortening (Lf) for SOL (D), GL (E), and VL (F); and mean normalised fascicle shortening velocity (Lf/s) for SOL (G), GL (H), and VL (I); and normalised fascicle to MTU shortening ratio for SOL (J), GL (K), and VL (L). * indicates a significant effect of grouping (p < 0.05).

### Joint Mechanics

There was a significant effect of grouping on mean knee moment, mean ankle and knee positive power, and ankle and knee positive work (all p < 0.001) but not mean ankle moment (p = 0.420) (Fig. 6). A significant hop frequency effect was found for mean knee moment (p < 0.001), mean knee positive power (p < 0.001), and ankle (p = 0.008) and knee (p < 0.001) positive work but not mean ankle positive power (p = 0.801). A significant hop height effect was found for mean ankle (p = 0.008) and knee (p < 0.001) positive power, and ankle (p = 0.008) and knee (p < 0.001) positive work but not mean knee moment (p > 0.999).

**Figure 6.**
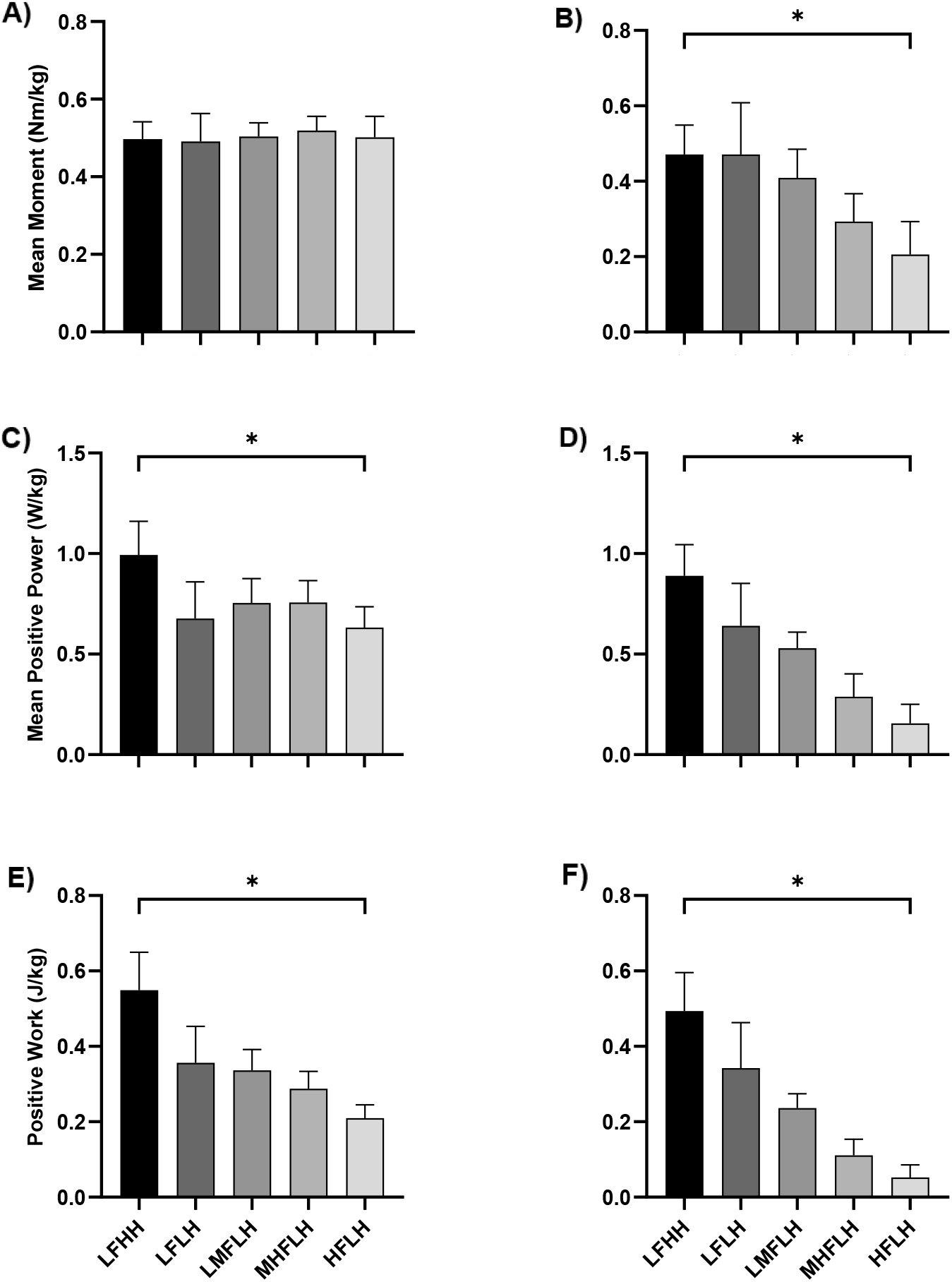
Group analysis of mean joint moment, mean positive joint power, and positive joint work data. The mean ± SD mean moment (Nm/kg) for each of the groups analysed for the ankle (A) and knee (B) joint; and mean positive work (J/kg) for the ankle (C) and knee (D); and mean positive power (W/kg) for the ankle (E) and knee (F). * indicates a significant effect of grouping (p < 0.05).

### Self-Selected Hopping

When completely unconstrained, subjects chose to hop to a mean height and frequency of 0.09 ± 0.02 m and 2.48 ± 0.24 Hz, respectively (Fig. 3a).

## Discussion

In this study, we sought to understand how different hopping conditions affect the apparent relationship between muscle mechanics and energy expenditure. To do so, we imposed a wide variety of frequency and height constraints to hopping and quantified gross metabolic power and the activation and work requirements of a range of key hopping muscles. Within the constraints that we imposed, we show that metabolic power increases with decreases in hop frequency and increases in hop height, and that increases in metabolic power are generally reflected by less economical muscle contractile mechanics.

As expected, increases in gross metabolic power were moderately associated with decreases in hop frequency and increases in hop height (Gutmann and Bertram, 2013). Hop frequency and height had a stronger association with gross metabolic power when used together in the regression model than when used singularly, indicating that hop frequency and height should be considered in combination when predicting the metabolic cost of hopping. However, these associations were still only moderate in strength, which is indicative of variability in the underlying muscle mechanics that drive metabolic cost across the cost landscape. Therefore, it was important to characterise how the requirements of key muscles change across a wide range of hopping conditions.

The mechanical requirements of ankle and knee musculature differed across conditions and were indicative of changes in metabolic power. The factors that are likely to contribute to increases in metabolic rate include increased average muscle force (i.e., activation requirements), and increased fascicle shortening and fascicle shortening velocity (i.e., work requirements) (Barclay, 2023; Homsher, 1972; Smith, 1972; Woledge et al., 1985). Below we explore potential contributors to changes in metabolic power with hop height and hop frequency.

At the ankle, the requirement to generate higher positive power and work, rather than average force, with increases in hop height or decreases in hop frequency likely contributes to increased metabolic power demands. There was no significant effect of grouping on average activation across the ankle muscles, with the exception of GM, athough this muscle did not have a clear frequency or height effect when tested with multiple comparisons. As such, ankle muscle activation costs will vary little with changes in hopping condition. SOL, GL and VL fascicle shortening and fascicle shortening velocity increased with decreased hop frequency, likely contributing to increased metabolic rate. While the net work done is similar across hop frequency conditions, there is clear requirement to do more positive work at lower hop frequencies (Fig. 6) and this is done primarily by muscles fibres rather than tendon, with a higher fascicle to MTU shortening ratio that likely contributes to increased metabolic costs (Dean and Kuo, 2011; Takeshita et al., 2006). Moreover, the counterintuitive increase in fascicle shortening velocity with reduction in movement frequency can be explained by the magnitude of the increase in fascicle shortening exceeding the relative increase in time to perform the movement. With increased hop height, only SOL fascicle shortening velocity showed a significant change. In contrast, the increase in fascicle shortening with increase in hop height likely results from reductions in duty factor (i.e., percentage of the hop cycle spent in contact with the ground), reducing the time by which fascicle length changes must occur during ground contact (Beck et al., 2020). The requirement to do more positive ankle work, at a higher average rate (i.e., power) with increased hop height, with no change in the contribution between muscle and tendon, was therefore largely attributed to SOL muscle function. Therefore, SOL likely contributes significantly to the increased metabolic rate with higher hop heights.

It is interesting to consider why mean EMG was unchanged about the ankle despite differences in the work requirements. This is probably a result of calculating mean EMG over the entire hop cycle, in contrast to quantifying work requirements only during the phase of positive CoM power. That is, more active musculature will be needed to produce an amount of force where positive work requirements are greater, but differences in the amount of eccentric work performed by muscle (Fig. S2), or in the overall duration of the hop cycle, or in duty factor could offset differences in mean EMG between conditions.

At the knee, work and power requirements closely reflected metabolic power changes across conditions, however this was partially driven by changes in force requirements. The increased activation of knee extensors (VL and RF) with decreased hop frequency stemmed from the requirement to generate increased force (i.e., knee moment). During hopping, knee flexion tends to increase as hop frequency decreases, increasing the GRF moment arm about the joint. This moment arm becomes larger about the knee than it does the ankle, resulting in higher force requirements, which helps to explain why differences in mean EMG were seen at the knee but not the ankle (Gutmann and Bertram, 2017a). Additionally, there was a significant increase in BF mean EMG with increased hop height, which might contribute to increased metabolic power with increased hop height, although it is expected that energy output of this muscle will likely be much lower than the extensor muscles. VL fascicle shortening and shortening velocity also increased concomitantly with increases in work and power at the knee as hop frequency decreased, which has the effect of increasing muscle energy rate (Lichtwark and Wilson, 2005b).

Within the constraints that we imposed, we found that metabolic power was positively related to hop height and negatively related to hop frequency, similar to Gutmann and Bertram (2013). However, had we forced subjects to hop to higher frequencies than we imposed, we may have eventually observed an increase in metabolic power that resembles the U-shaped relationship classically seen between movement frequency and metabolic cost, assuming a constant height. Studies have shown that hopping to frequencies upwards of 3.0 Hz can begin to increase active muscle work as a result of muscle operating at shorter than optimal lengths and acting out of phase with tendon (Dean and Kuo, 2011; Monte et al., 2021; Robertson and Sawicki, 2014, 2015).

Notably, self-selected hopping did not minimise metabolic power on average (Fig. 3b), despite minimisation of metabolic cost being widely regarded as a principle optimality criterion governing locomotion (Brown et al., 2021; Gutmann and Bertram, 2013; Ralston, 1958; Sellinger et al., 2015). Certainly, the minimisation of metabolic cost may make up part of this control scheme, although, in our case this might be secondary to other factors such as the minimisation of muscle activation (McDonald et al., 2022; McMahon et al., 1987; Miller et al., 2012; Neptune and Hull, 1999). Self-selected hopping did not minimise metabolic power because it would not have minimised the work requirements of the power generating muscles. Had participants elected to hop to higher frequencies on average (i.e., greater than 2.48 Hz) this would likely have led to more isometric muscle behaviour.

We believe our results are important for informing biomechanical models of the metabolic cost of hopping and other forms of bouncing locomotion. Our results seem to echo the work of Gutmann and Bertram (2017a) and Allen et al. (2022), who suggest that knee muscle mechanics, and proxies thereof, should be especially indicative of changes in energy expenditure during hopping since knee musculature dominates changes in total active muscle volume across hopping conditions. However, this may not entirely translate to other bouncing gaits such as running, where, especially at higher step frequencies, the percentage contribution of the knee to changes in total active muscle volume can become less of that of the ankle and hip (Kipp et al., 2018). This is likely driven by the fact that as we run at higher speeds and frequency of strides, there is also a change in stride length requiring increased flight time and height per stride (Hunter et al., 2004), which our results suggests requires increased ankle power.

There are limitations to this study that should be considered. Though the mechanics and energetics of hopping are at the core of other bouncing gaits, hopping omits features of other gaits (e.g., leg and arm swing) that influence their metabolic cost. Moreover, as mentioned previously, the contributions of different joints to changes in total active muscle volume, and therefore whole-body metabolic cost, may not vary the same way for hopping as they do for other bouncing gaits. Furthermore, we acknowledge that properties of muscle and tendon (e.g., fascicle lengths, muscle volumes, tendon stiffnesses) can vary substanitally between individuals and thereby vary the mechanics and energetics elicited by different hopping conditions. Whilst effects of participant were factored into our analyses, it is still possible that our findings are subject to a degree of interindividual variability. Lastly, whilst this study provides an indication of how the activation and work requirements of different muscles change alongside changes in metabolic power, the results do not allow us to partition the contributions of muscle activation and work to metabolic power, and we were limited to data from select muscles. We suggest that musculoskeletal modelling approaches, validated against such datasets, may provide more information than can be garnered by in-vivo measures alone.

In conclusion, within the constraints that we imposed, reductions in hop frequency and increases in hop height resulted in increases in metabolic power that could be explained by increases in the activation requirements of knee musculature and/or increases in the work requirements of both knee and ankle musculature. To help direct future biomechanical models of energy expenditure, we hope that our approach or similar can be extended to other gaits to characterise potential variance in muscle mechanics and thus metabolic energy expenditure.

## Authorship Statement

**LN Jessup:** Conceptualisation, Methodology, Recruitment, Data collection, Data processing, Writing – original draft, Writing – review & editing, Visualisation. **LA Kelly:** Conceptualisation, Methodology, Writing – review & editing, Supervision. **AG Cresswell:** Methodology, Writing – review & editing, Supervision. **GA Lichtwark:** Conceptualisation, Methodology, Data processing, Writing – review & editing, Supervision.

## Acknowledgements

We thank all subjects, who each volunteered many hours of their time to participate in this study.

## Competing Interests

No competing interests declared.

## Funding

LN Jessup is supported by a University of Queensland Graduate Student Scholarship. GA Lichtwark and research costs are supported by an Australian Research Council Future Fellowship (FT190100129). LA Kelly is supported by an Australian Research Council DECRA Fellowship (DE200100585).

## Notes

### Competing Interest Statement

The authors have declared no competing interest.

